# Quantitative measurement of activity of JAK-STAT signaling pathways in blood samples and immune cells to predict innate and adaptive cellular immune response to viral infection and accelerate vaccine development

**DOI:** 10.1101/2020.05.13.092759

**Authors:** Wilbert Bouwman, Wim Verhaegh, Laurent Holtzer, Anja van de Stolpe

## Abstract

The host immune response determines the clinical course of a viral infection, for example in case of COVID-19 infection. The effectiveness of vaccination also depends on the induced immune response. Currently there is no method to measure the cellular immune response in blood samples. The functional activity of cells of innate and adaptive immune system is determined by coordinated activity of signaling pathways, especially the JAK-STAT pathways. Using a previously described approach we developed mRNA-based tests to measure activity of these signaling pathways, and show that they can be used to measure in a quantitative manner the cellular innate and adaptive immune response to a viral infection or vaccine in whole blood, PBMC, and specific immune cell type samples. Pathway activity level and range in healthy individuals was established, enabling interpretation of a pathway activity score on a patient sample without the need for a reference sample. Evidence is presented that the pathway activity analysis may also be useful for in vitro vaccine development and assessment of vaccine immunogenicity. Other envisioned applications lie in development of immunomodulatory drugs and drug response prediction and monitoring. Tests are expected to be of value in the COVID-19 crisis. In addition to the described Affymetrix microarray-based pathway tests for measuring host immune response, qPCR-based versions are in development; the latter can in principle be performed within three hours in routine hospital labs.

## Introduction

A good functioning immune system is crucial to protect against diseases, such as viral and bacterial infections and cancer (1). It generates an immune response to non-self antigens, such as a component of a virus, or an abnormal protein produced by a cancer cell. Briefly and highly simplified, an immune response consists of an innate immune response which has an inflammatory component and is responsible for rapid non-specific attack and removal of pathogens and for presentation of pathogen-derived antigens to the cells of the adaptive immune response, which is responsible for generating long term immunity. Both parts of the immune system contain different immune cell types that cooperate to generate an effective immune response. They communicate via a large number of cytokines, such as interleukins and interferons. Cytokines convey messages from one cell to the other by binding to specific cellular receptors to activate cellular signal transduction pathways, especially the JAK-STAT pathways, resulting in production of proteins that adapt the function of the immune cell as response to the cytokine signal. In this highly organized system, dendritic cells are the most important antigen presenting cells bridging the innate and adaptive immune system. The adaptive immune response consists of a humoral response, that is, antibodies specifically generated against the pathogen by B-cells, and a cellular immune response, effectuated by cytotoxic T-cells. For an adequate response to a dominantly intracellular pathogen, such as a virus or in case of cancer cells, the cellular immune response is of prime importance, in addition to the production of neutralizing antibodies. The effectiveness with which the immune system can mount a response to a pathogen is determined by multiple factors, such as the amount and type of pathogen, genetic variations in relevant immune genes, patient comorbidities, and use of drugs which may have a suppressive, or stimulating, effect on the immune system.

Viral infections can create worldwide disasters, as exemplified by the COVID-19 pandemic, caused by the SARS-CoV-2 virus (2). A major clinical problem caused by infection with this respiratory virus is rapid deterioration in a subset of patients with pneumonia, requiring admission to an intensive care unit (ICU), frequently with prolonged invasive ventilation and detrimental outcome (3),(4). Other respiratory viruses, such as other coronaviruses (e.g. SARS and MERS) but also other viruses like the Respiratory Syncytial Virus (RSV) can cause similar clinical behavior in a subset of patients. Many other types of viruses, that do or do not specifically target the respiratory system, can also cause epidemics associated with high mortality and serious complications, e.g. influenza, ebola, measles, yellow fever, etc.

The clinical course of a viral infection is strongly determined by the host immune response, emphasized in the case of COVID-19 by the extreme differences in symptomatology and clinical outcome, ranging from mild symptoms, or even unnoticed infection, to Acute Respiratory Distress Syndrome (ARDS) with high mortality rate (2),(3),(5). Measuring the functional state of the innate and adaptive immune response in a blood sample of a patient is expected to improve prognosis prediction, and to support development of vaccines and safe immunomodulatory treatments to improve immune response and clinical outcome.

We describe development, as well as evaluation in preclinical and clinical studies, of an mRNA-based test to quantify the immune response to a viral infection, based on measuring activity of the JAK-STAT1/2 and JAK-STAT3 signal transduction pathways in a whole blood or Peripheral Blood Mononuclear Cell (PBMC) sample, or in separate immune cell types. The technology used to develop tests to measure activity of signal transduction pathways is based on measuring the mRNA levels of target genes of the pathway-specific transcription factor and has been described before (6),(7),(8),(9),(10),(11),(12),(13).

## Methods

### Development of JAK-STAT1/2 and JAK-STAT3 pathway tests

Development of tests to measure JAK-STAT1/2 and JAK-STAT3 pathway activity were developed as described before (6),(7),(8). In brief, target genes of the STAT transcription factors of the respective pathways were identified, and a Bayesian network computational model was created for interpretation of measured mRNA levels of the pathway target genes. For the purpose of this study, target gene mRNA levels were derived from Affymetrix expression microarray data (6).

The mathematical approach to develop Bayesian network models for the interpretation of the transcription factor target gene mRNA levels has been described previously in detail (6). A causal computational network model is built, enabling inference of the probability that the pathway-associated transcription factor is in an active state, binding to and transcribing its target genes (Figure S1). The Bayesian network describes (i) the causal relation that a target gene is up- or downregulated depending on the transcription factor being active or inactive and (ii) the causal relation that an AffymetrixU133Plus2.0 probeset signal is high or low, depending on the target gene being up- or down regulated. Model parameters for (i) have been based on literature evidence, and for (ii) are based on data from calibration samples with known signalling pathway activity, that is, either active or inactive. Target genes for the JAK-STAT1/2 and JAK-STAT3 pathway model were selected using available scientific literature (Supplementary Table S2, and Supplementary methods). Following model calibration and model parameter freeze, mRNA probeset measurements of a new sample can be entered into the model, and Bayesian inference is used to calculate probability *P* that the transcription factor, and therefore the signalling pathway, is active. Pathway activity score can be presented as log2odds value log2 (*P* / (1 – *P*)), which can optionally be normalized on a scale from 0-100 (6),(7),(8). Quantitative log2odds pathway activity scores are used in the figures and results description. JAK-STAT1/2 and JAK-STAT3 Pathway models were biologically validated on independent public available (14) Affymetrix HG-U133Plus2.0 microarray data of samples with known pathway activity.

### Analysis of Affymetrix microarray datasets from preclinical and clinical studies

Using the developed signaling pathway tests for JAK-STAT1/2 and JAK-STAT3 pathways, public Affymetrix HG-U133Plus2.0 microarray expression datasets from preclinical and clinical studies (deposited in the GEO database (14)) were analyzed and log2odds pathway activity scores calculated. In one dataset a previously described pathway test to measure NFκB pathway activity was used (8).

### Microarray data quality control

Quality control (QC) was performed on Affymetrix data of each individual sample (15),(16). In summary, QC parameters include: the average value of all probe intensities, presence of negative or extremely high (> 16-bit) intensity values, poly-A RNA (sample preparation spike-ins) and labelled cRNA (hybridization spike ins) controls, *GAPDH* and *ACTB* 3’/5’ ratio, centre of intensity and values of positive and negative border controls determined by affyQCReport package in R, and an RNA degradation value determined by the AffyRNAdeg function from the Affymetrix package in R (17),(18). Samples that failed QC were removed prior to data analysis.

### Statistics

Boxplots and individual sample plots were made using the Python data visualization library function *seaborn*; additional statistical annotations were created using the Python package statannot (19),(20). T-test or two sided Mann-Whitney-Wilcoxon testing was used to compare pathway activity scores across groups. p-values are indicated in the figures.

## Results

### Development and validation of the JAK-STAT1/2 and JAK-STAT3 pathway assay

Target genes that were identified and used to develop the JAK-STAT1/2 models and the JAK-STAT3 models are listed in Supplementary Table S1. The Bayesian model developed for the JAK-STAT1/2 pathway was calibrated on Affymetrix data from samples stimulated with IFN type II (GSE38351, (21)). Both types of interferon activate the JAK-STAT1/2 pathway (22). In this study, PBMC samples from patients with rheumatoid arthritis had been stimulated with type I and type II IFN, while untreated samples and TNFα-stimulated samples were used as STAT1/2 pathway inactive controls (Figure S2A). The calibrated JAK-STAT1/2 pathway model measured both IFN type I and type II activity of the JAK-STAT1/2 pathway (Figure S2B). Biological validation of the calibrated and frozen pathway model was performed on independent datasets in which either IFN type I or IFN type II had activated the pathway. Pathway activity was correctly measured in natural killer (NK) cells stimulated *in vitro* with IFNα-2b (IFN type I) (Figure S2C) (GSE15743, (23)), and in dendritic cells stimulated with IFNβ (IFN type I) (Figure S2D) (GSE52081, (24)) Stimulation with IFNγ (IFN type II) induced an increase in pathway activity scores in respectively THP-1 monocytic cells (Figure S2E) (GSE58096, (25)), dendritic cells (Figure S2F) (GSE11327, (26)), and monocytes (Figure S2G) (GSE11864, (27)).

The JAK-STAT3 pathway can be activated by several cytokines that bind to their respective receptors, resulting in activation of JAK1 and formation of a STAT3 homodimer which transcribes target genes (28),(29). The JAK-STAT3 pathway model was calibrated on T cell leukemia cells, in the absence or presence of interleukin 2 (IL2) stimulation to activate the pathway (Figure S3A) (GSE8687, (30)). Validation of the JAK-STAT3 pathway model was performed on independent datasets with known JAK-STAT3 pathway activity: IL-2, IL-15, and IL10, known to induce JAK-STAT3 pathway activity, induced an increase in pathway activity scores in a Sezary (Sez) cutaneous lymphoma cell line and in PBMCs (Figures S3B, S3C) (GSE8685, (30); GSE43700, (31)). IgG was also reported to induce JAK-STAT pathway activity and incubation of the Sez cell line with IgG-coated beads increased JAK-STAT3 pathway activity scores (Figure S3D) (GSE8507, (32)). In clinical patient sample data of knee synovial biopsies from patients with rheumatoid arthritis JAK-STAT3 pathway activity scores decreased after treatment with Tocilizumab, an inhibitor of the IL6 ligand of the JAK-STAT3 pathway, and not after treatment with methotrexate (Figure S3E) (GSE45867, (33)). In summary, the JAK-STAT1/2 and the JAK-STAT3 pathway model performed as expected on independent preclinical and clinical sample data.

### Measuring JAK-STAT1/2 and JAK-STAT3 pathway activity in blood samples of clinical viral infection studies

JAK-STAT signaling pathways transduce cytokine signals induced by a viral infection into functional changes in immune cells. To investigate potential clinical value of measuring JAK-STAT1/2 and JAK-STAT3 pathway activity in patients with a viral infection, pathway activity was measured on gene expression data from clinical studies on infections with influenza, respiratory syncytial virus) (RSV), dengue, yellow fever, rotavirus, and hepatitis B virus.

#### Activity of the JAK-STAT1/2 pathway in PBMCS distinguished between viral and bacterial infectious disease

In a clinical study in which PBMC sample data were analyzed from patients with an influenza virus infection and from patients with a bacterial infection with either S. Pneumonia or S. Aureus, activity of the JAK-STAT1/2 pathway was significantly increased in patients with a viral infection. In contrast, activity of the JAK-STAT3 pathway did not distinguish sufficiently between viral and bacterial infections (Figure 1).

**Figure 1.**
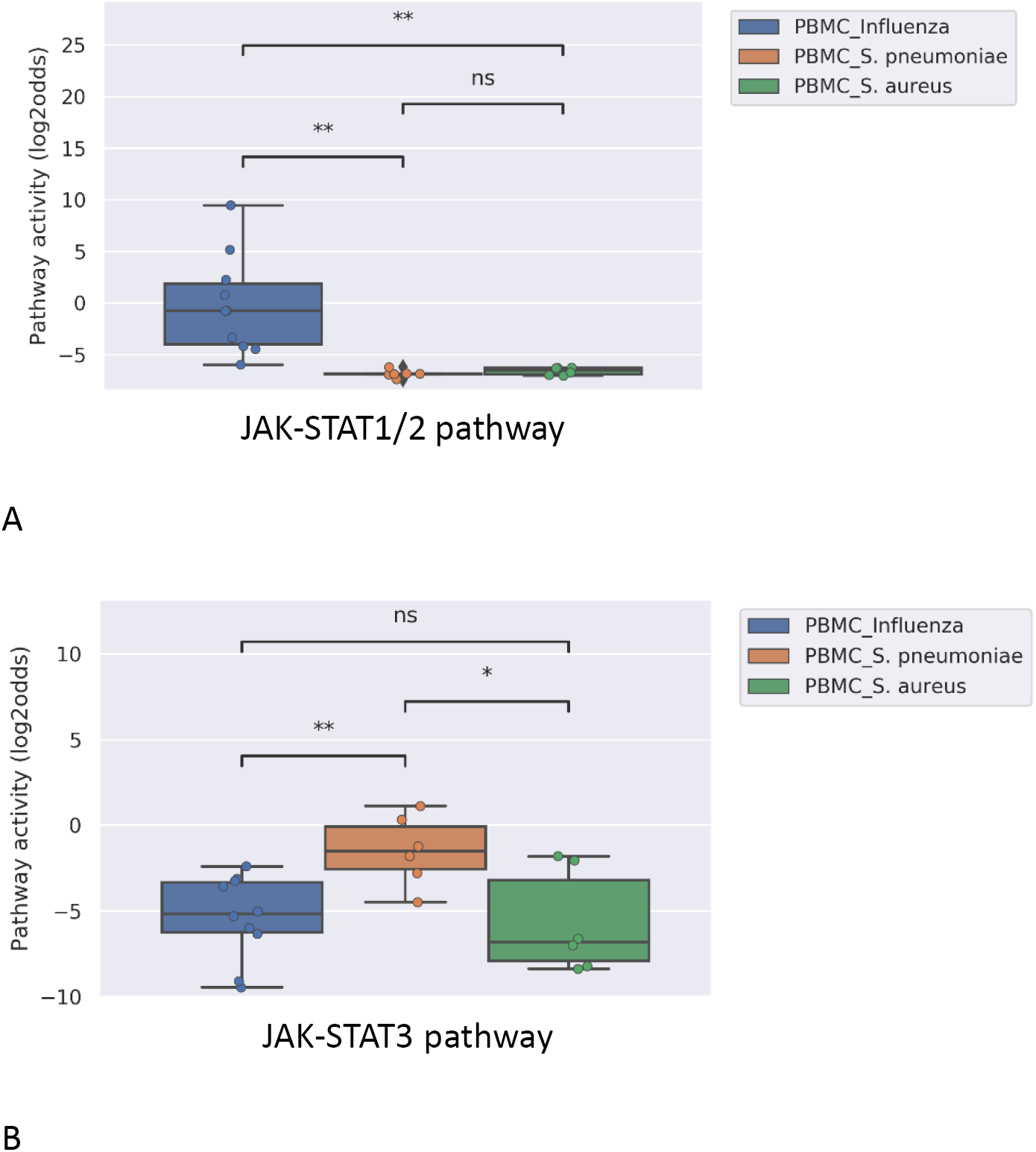
JAK-STAT1/2 (A) and STAT3 (B) pathway activity analysis of dataset GSE626. PBMC samples of pediatric patients with acute infection with influenza virus (blue), S. Pneumoniae (orange) or S. Aureus (green). Pathway activity score on Y-axis in log2odds. Two sided t-test independent tests were performed; p-values are indicated in the figures. P-values indicate: *p<0.05; **p<0.01.

#### JAK-STAT1/2 and JAK-STAT3 pathway activity are induced in blood cells by viral infection and JAK-STAT3 pathway activity distinguished between mild and severe infection

Some viral infections are known for their variation in disease severity. The RSV virus is a respiratory RNA virus which predominantly infects children and presents with a large variation in disease severity. Dengue virus infection also can run a more severe disease course, presenting as Dengue Hemorrhagic Fever (DHF). In PBMC samples of patients with one of these viral infections, JAK-STAT1/2 pathway activity scores were increased irrespective of the degree of severity of the infection (Figure 2A and 3A). However, activity of the JAK-STAT3 pathway was significantly higher in RSV patients who presented with a severe disease course, and also higher in patients with DHF, however the latter did not reach significance (n=3) (Figure 2B and 3B). In a separate clinical study, CD8+ T cells had been isolated from patients with Dengue and Dengue Hemorrhagic Fever (DHF). In this T cell subtype, again activity of both JAK-STAT pathways was increased in patients, but JAK-STAT3 pathway activity was not higher in the severe DHF disease variant (Figure 3C,D). Dengue virus can induce partial immunosuppression, and the PBMC blood samples that had been collected at various time points during the disease enabled investigation of the relation between JAK-STAT pathway activity and this immunosuppressive effect of the virus. JAK-STAT1/2 pathway activity rapidly decreased; in DHF the decrease was already significant two days after clinical presentation, in DF after day 5, while the JAK-STAT3 pathway activity decreased more slowly during the disease (Figure 3A,B).

**Figure 2:**
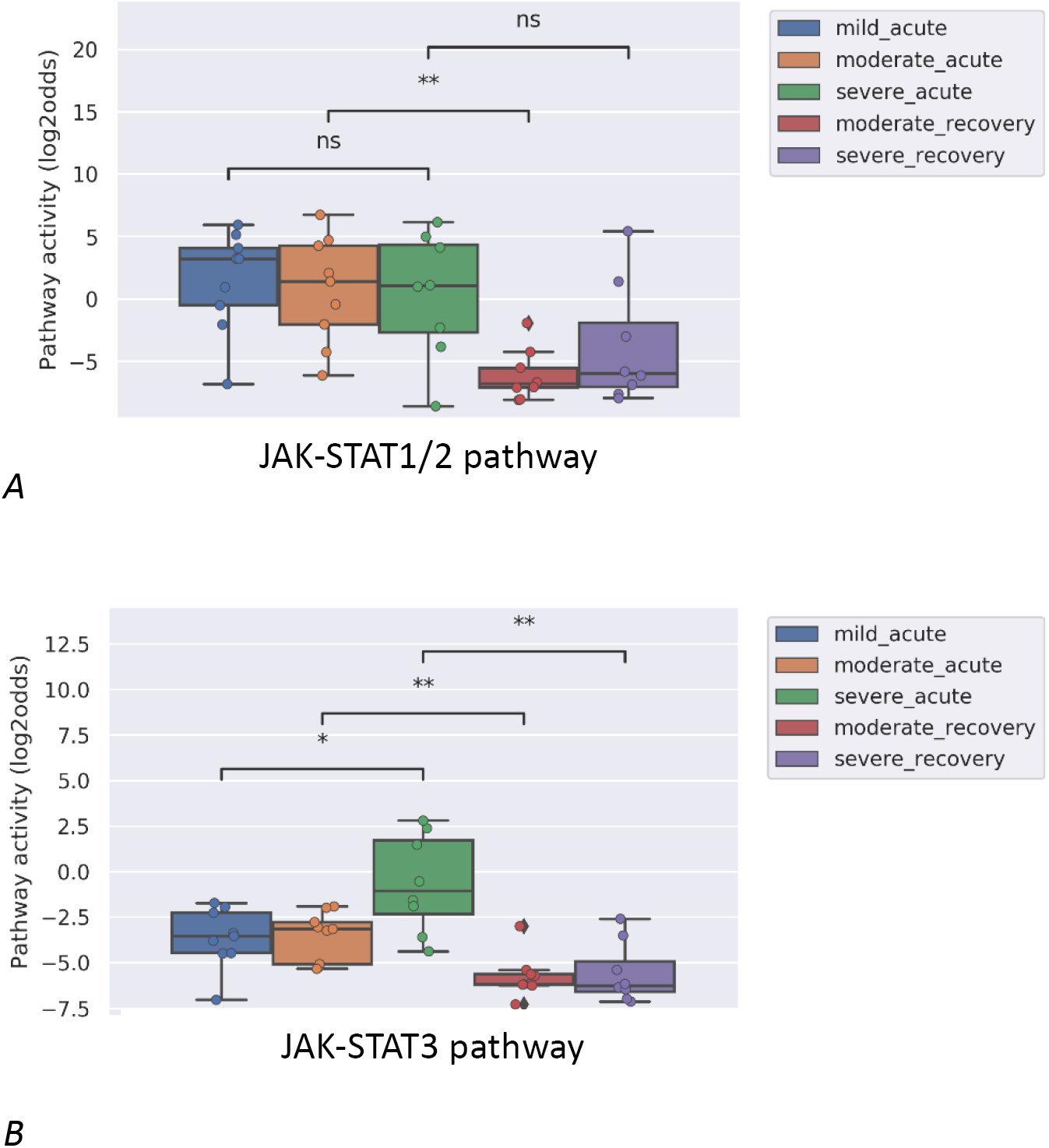
JAK-STAT1/2 (A) and JAK-STAT3 (B) pathway activity analysis of dataset GSE69606 (34) including patients with acute RSV infections, divided into mild, moderate and severe disease. From moderate and severe diseased patients recovery samples (on average 4 weeks after discharge) were obtained. Peripheral blood mononuclear cells (PBMCs) were used for analysis. Pathway activity score on Y-axis in log2odds. Two sided Mann–Whitney–Wilcoxon statistical tests were performed; p-values are indicated in the figures. P-values indicate: *p<0.05; **p<0.01.

**Figure 3:**
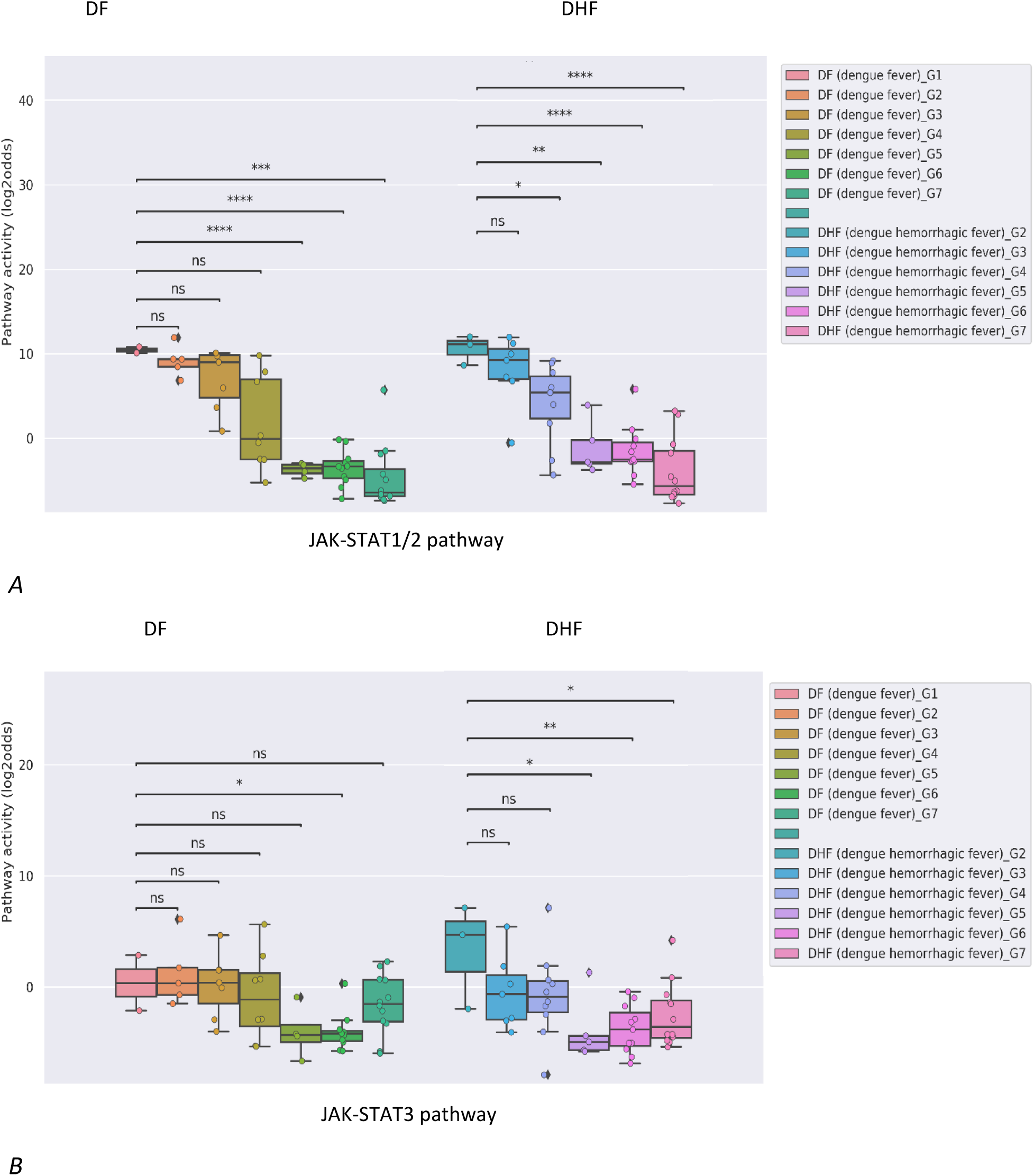
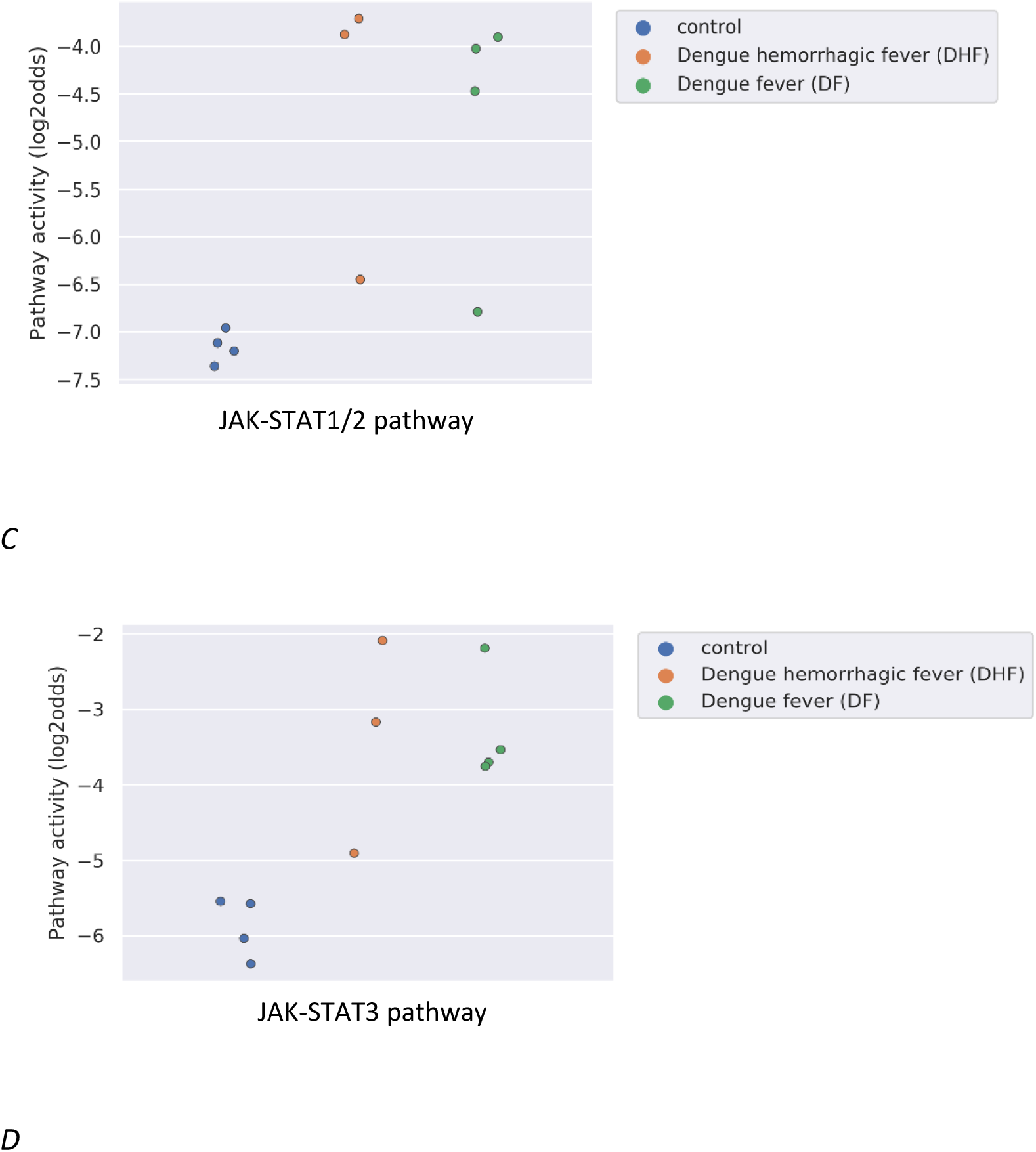
JAK-STAT1/2 and STAT3 pathway activity analysis of dataset GSE43777 (A for STAT1-2 and B for STAT3) and dataset GSE84331 (C for STAT1-2 and D for STAT3). 3A-B. Dataset GSE43777 (35) including DF (dengue fever) and DHF (dengue hemorrhagic fever) patient samples during the course of dengue acute illness. Analysis was performed on PBMCs. Each stage represents a group of patients after x days from fever onset, Stages (days after onset fever): G1: 0 days, G2: 2 days, G3: 3 days, G4: 4 days, G5: 5 days, G6: 6-10 days, G7: >20 days. 3C-D. Dataset GSE84331 (36) includes CD8+ T cells (HLA-DR+CD38+) from Dengue/DHF patients which were compared to naive (CCR7+CD45RA+) CD8+ T cells from healthy donors (Control). Pathway activity score on Y-axis in log2odds. Two sided t-test independent statistical tests were performed; p-values are indicated in the figures. P-values indicate: *p<0.05; **p<0.01.

#### JAK-STAT1/2 pathway activity in PBMC samples reflected the strength of induced immunity by a specific virus type

Influenza virus is known to generally induce a stronger immunity than RSV. In the analyzed study, this was associated with higher JAK-STAT1/2 pathway activity, but not JAK-STAT3 pathway activity, in PBMCs of patients with influenza infection, compared to RSV infected patients (Figure 4).

**Figure 4:**
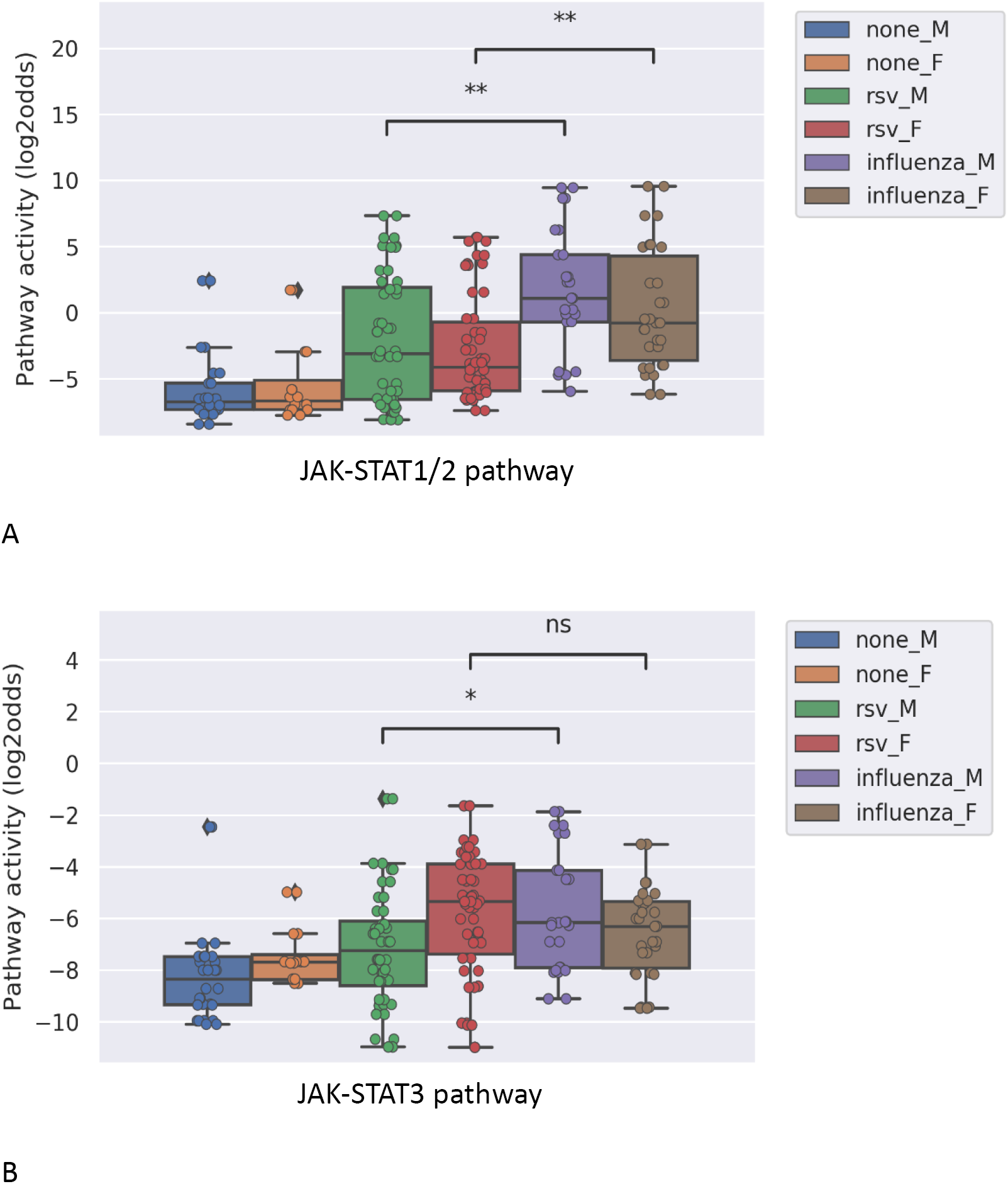
JAK-STAT1/2 (A) and STAT3 (B) pathway activity analysis of dataset GSE34205 (37), including children (M=male; F=female) of median age of 2.4 months (range 1.5-8.6) hospitalized with acute RSV and influenza virus infection. Blood samples were collected within 42-72 hours of hospitalization and PBMCs were isolated for analysis and compared with healthy controls. Pathway activity score on Y-axis in log2odds. Two sided Mann–Whitney–Wilcoxon statistical tests were performed; p-values are indicated in the figures. P-values indicate: *p<0.05; **p<0.01; ***p<0.001.

#### Lack of increase in JAK-STAT1/2 pathway activity and reduced JAK-STAT3 pathway activity in dendritic cells in chronic Hepatitis B infection

Hepatitis B virus (HBV) may cause chronic liver infection with hepatitis, ultimately leading to severe complications such as liver cirrhosis and cancer. Plasmacytoid dendritic cells (pDCs) from blood of both asymptomatic HBV carriers and chronic HBV hepatitis patients, JAK-STAT1/2 pathway activity was not increased compared to healthy individuals, and JAK-STAT3 pathway activity scores were even lower than normal (Figure 5A,B).

**Figure 5:**
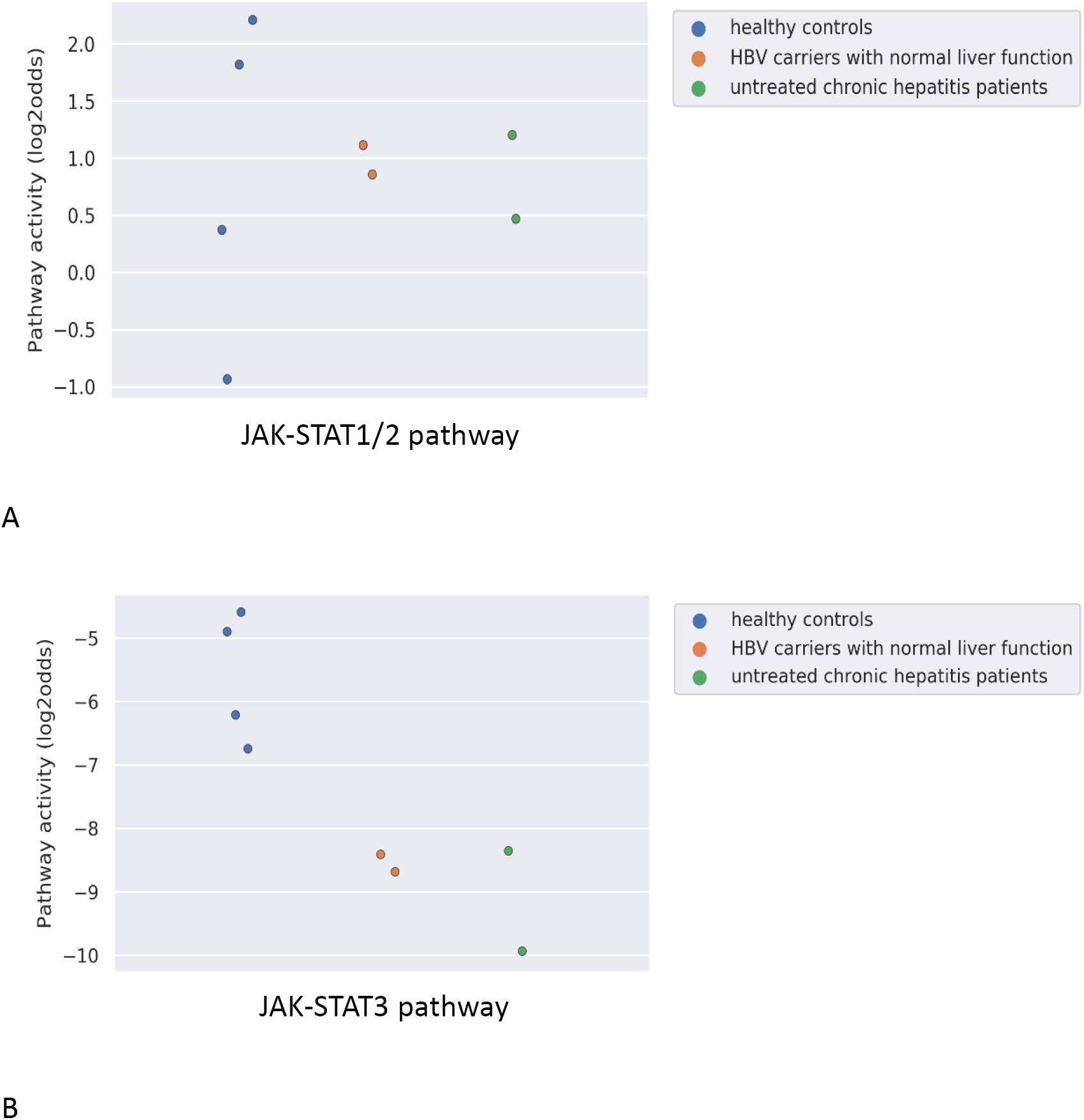
JAK-STAT1/2 (A) and STAT3 (B) pathway activity analysis of dataset GSE119322 (38) including HBV patients with normal liver function, untreated chronic hepatitis patients and healthy controls. Mononuclear cells were isolated and the plasmacytoid DCs (pDCs) were used for analysis. Pathway activity score on Y-axis in log2odds.

#### Measuring the effectiveness of vaccination in humans

Three clinical vaccination studies with yellow fever and influenza virus were analyzed. For yellow fever vaccination, healthy individuals were immunized with a live attenuated strain of the yellow fever virus (YFV-17D). In PBMCs (Figure 6A), as well as in CD4+ T cells and two monocyte subsets (Figure 6C, isolated before and after vaccination with the *live* Yellow Fever vaccine, JAK-STAT1/2 pathway activity scores increased after vaccination, compatible with induction of immunity (Figure 6A,C). JAK-STAT3 pathway activity scores were only increased in monocytes (Figure 6D). In contrast to Yellow Fever vaccination, vaccination with Trivalent *inactivated* nfluenza Vaccine (TIV) did not induce an increase in JAK-STAT1/2 or JAK-STAT3 pathway activity scores in PBMCs within the same time frame (one week) after vaccination, suggesting that this vaccination had not induced full immunity (Figure 6E,F).

**Figure 6:**
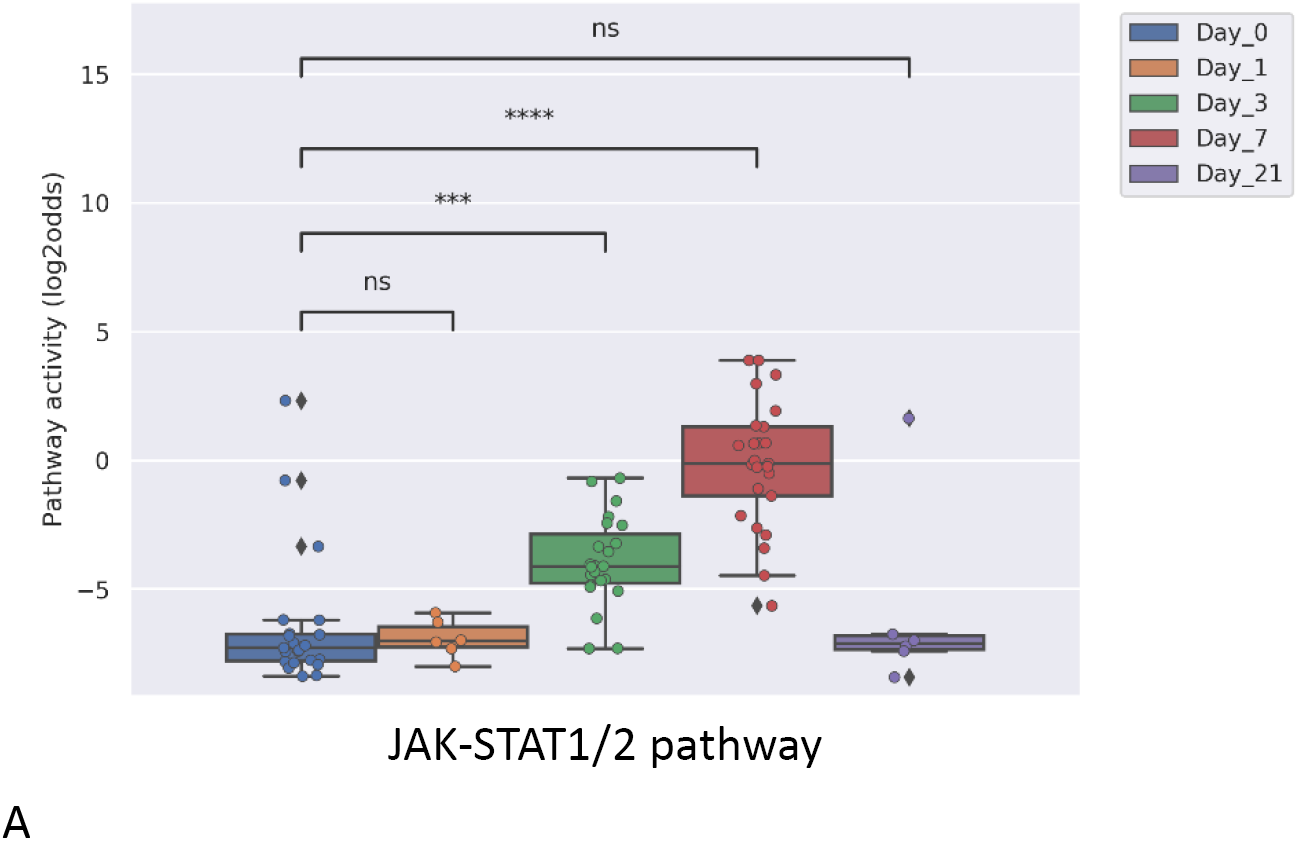
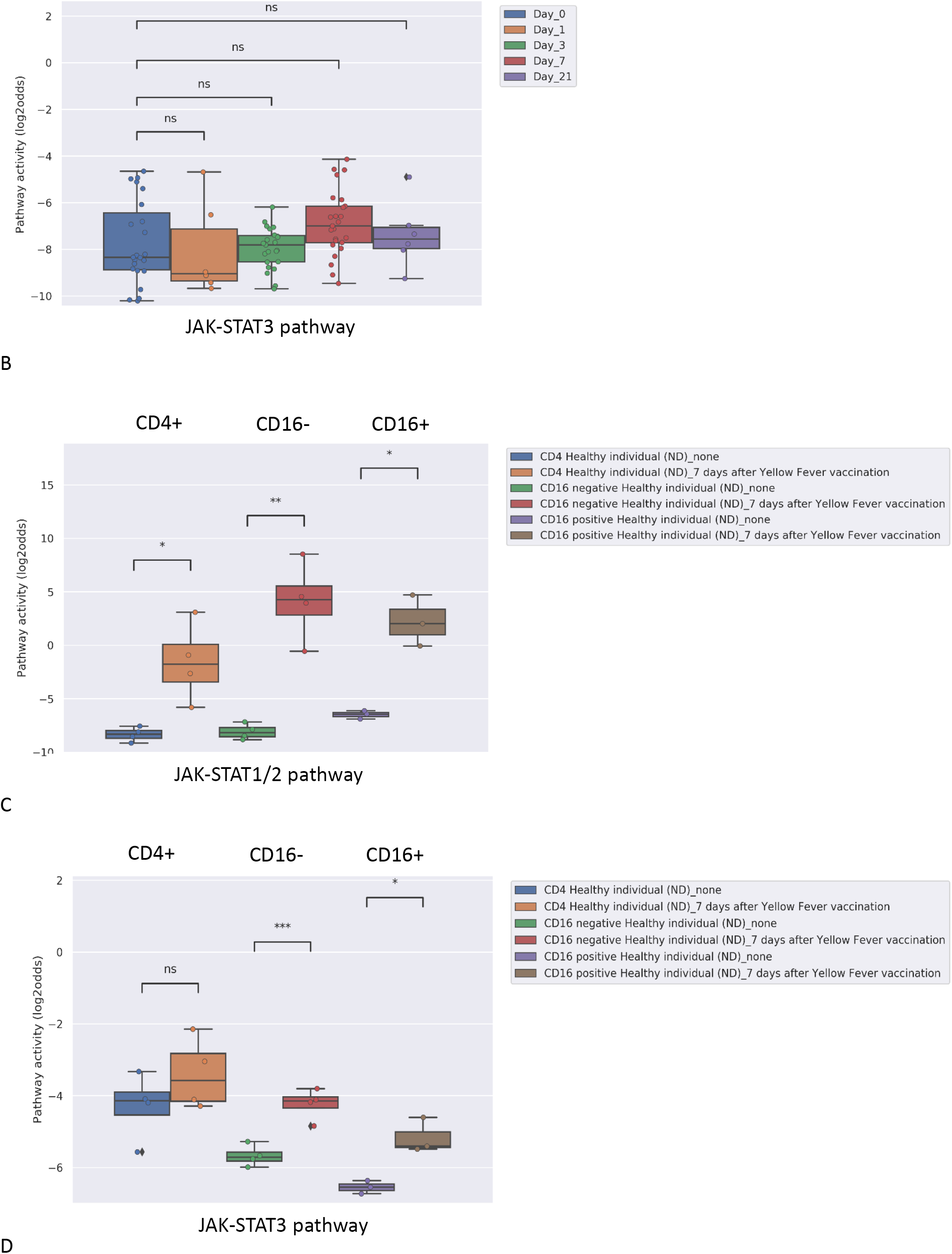
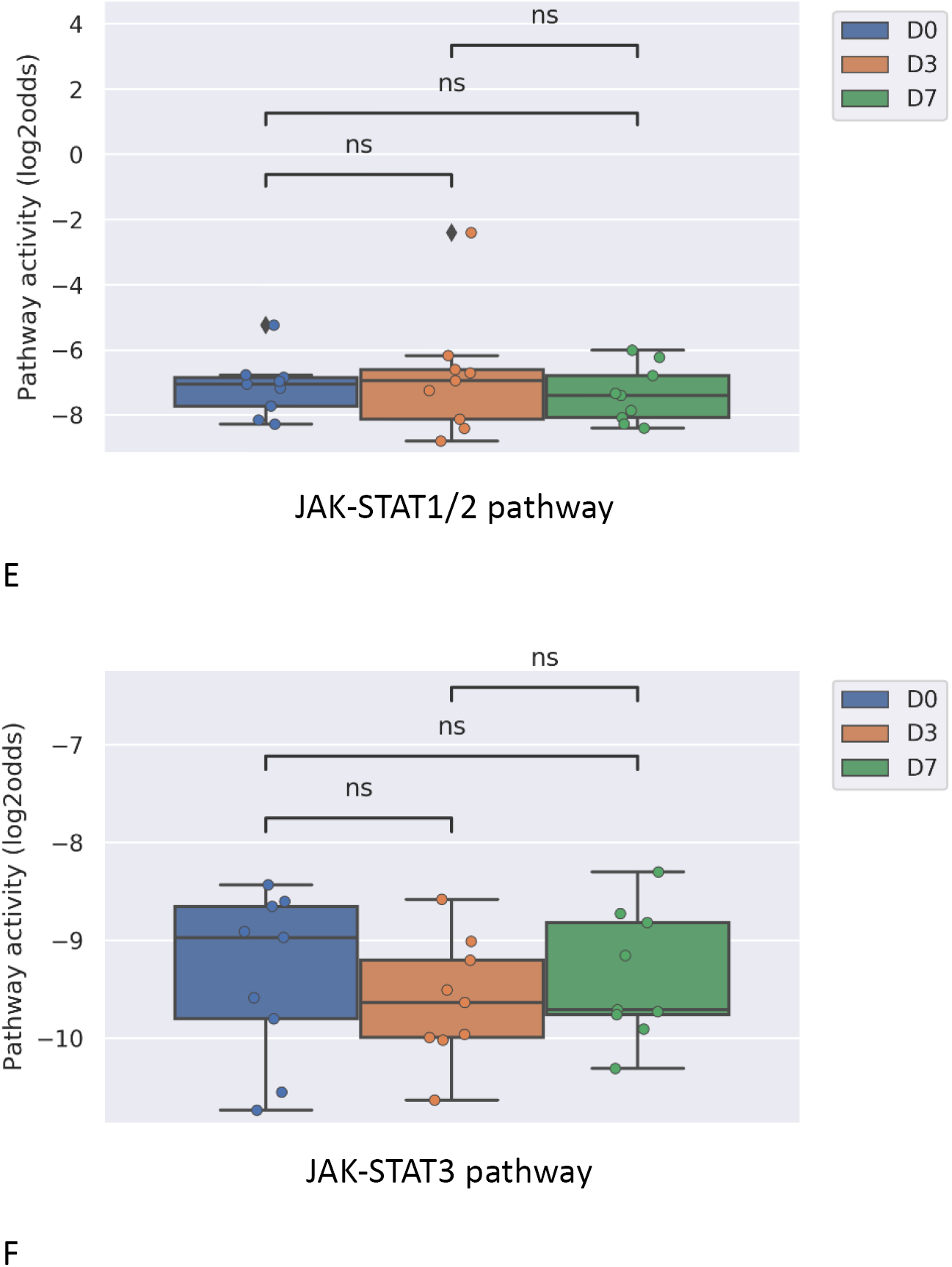
JAK-STAT pathway activity induced by vaccination in healthy individuals with Yellow Fever vaccine (YFV-17, live attenuated yellow fever virus strain without adjuvant) (A-D) or influenza vaccine (TIV, Trivalent Influenza Vaccine without adjuvant)(E,F). A,B: JAK-STAT1/2 (A) and JAK-STAT3 (B) pathway activity analysis of dataset GSE13486 (39). Yellow Fever vaccinated (YF-17D) healthy individuals at days 0, 1, 3, 7 and 21 after vaccination. PBMCs were isolated from blood and were used for analysis. C,D: JAK-STAT1/2 (C) and JAK-STAT3 (D) pathway activity analysis of dataset GSE51997 (21). Yellow fever vaccinated female volunteers. Peripheral blood was taken 7 days after immunization, and CD4 positive T-cells, CD16 negative and positive monocytes were isolated and used for analysis. E,F: JAK-STAT1/2 (E) and JAK-STAT3 (F) pathway activity analysis of dataset GSE29614 (40). Young adults vaccinated with Influenza TIV (Trivalent Inactivated Influenza Vaccine, GSK) vaccine during 2007/08 Flu Season. PBMC samples were isolated at days 0, 3, 7 post-vaccination. Pathway activity score on Y-axis in log2odds. Two sided t-test independent statistical tests were performed; p-values are indicated in the figures. P-values indicate: *p<0.05; **p<0.01; ***p<0.001; ****p<0.0001.

#### Measuring viral immunogenicity in DCs in vitro

Dendritic cells are the most powerful antigen presenting cells, bridging the innate immune response to the adaptive immune response. Both maturation and activation of DCs is required to enable processing and presentation of the viral antigen within the HLA complex for efficient T cell activation, requiring activation of the NFκB and JAK-STAT1/2 signal transduction pathways (41),(42). In the here analyzed study, dendritic cells had been infected in vitro with two HIV viral vectors separately or in combination. Only the combined infection with the two vectors was reported to be competent in fully activating the DCs, but neither of them alone. Pathway analysis showed that only in case both virions were present pathway activity scores of NFκB and JAK-STAT1/2 signaling pathways increased (Figure 7A,B).

**Figure 7:**
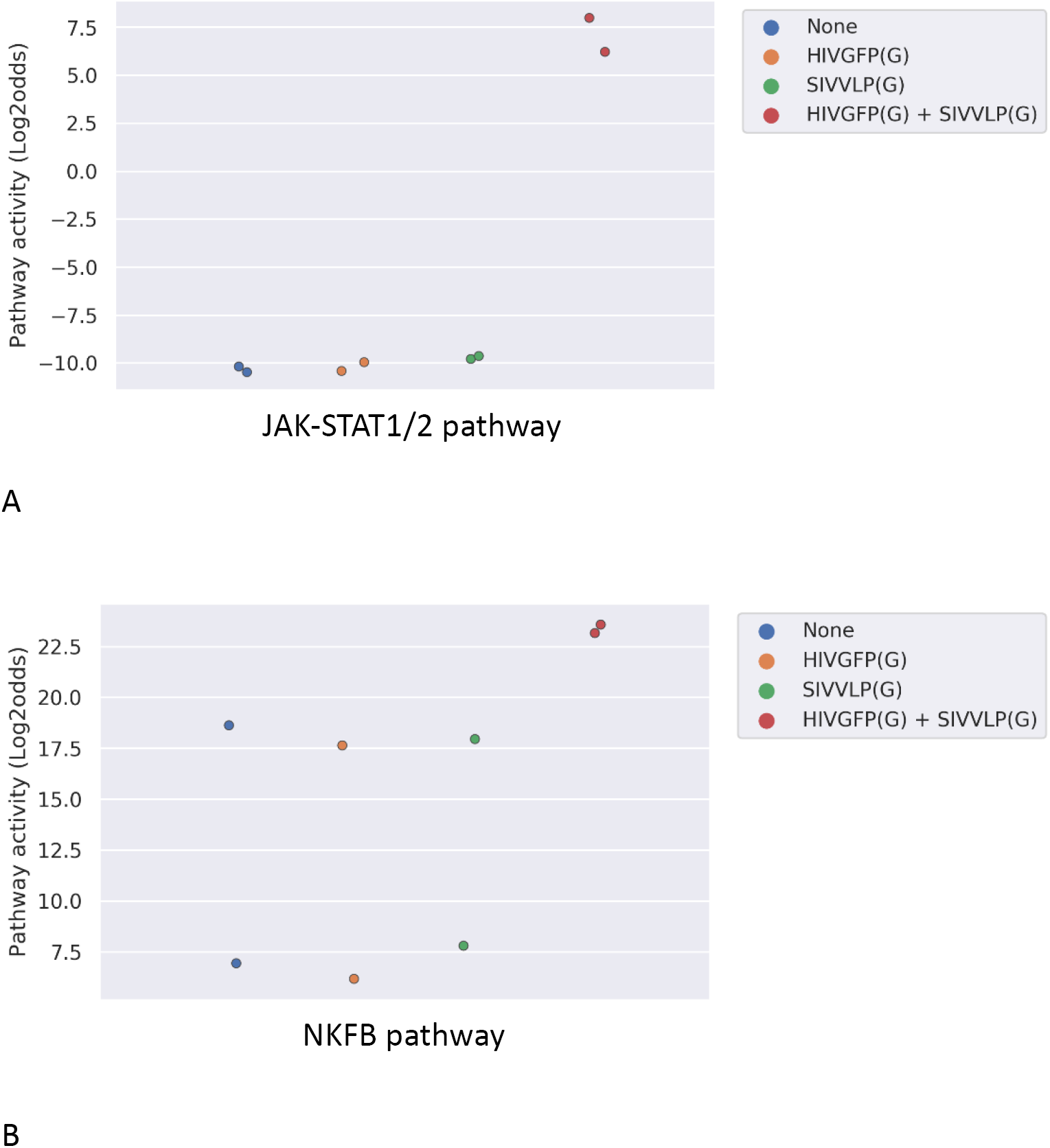
JAK-STAT1/2 (A) and NFκB (B) pathway activity analysis of dataset GSE22589 (n=2). Monocyte-derived dendritic cells (MDDCs) infected with an envelope-defective GFP-encoding VSV-G-pseudotyped HIV-1 vector (HIVGFP(G)) and with VSV-G pseudotyped virus-like particles derived from SIVmac to deliver Vpx (SIVVLP(G)), alone or in combination. Cells were infected with one or combination of the vectors at day 4 of differentiation and were harvested 48 hours later. Pathway activity score on Y-axis in log2odds.

#### Measuring cellular host immune response on whole blood samples of patients with a viral infection

In clinical practice it is expected to be of value to have a rapid and cost-effective test to diagnose a viral infection and measure viral infection- or vaccine-induced host immune response. A whole blood sample is easiest and least expensive to analyze in a routine clinical setting, but contains a mixture of cell types. Analysis of a clinical study in which whole blood samples (collected in PAXgene tubes) were obtained from patients with acute influenza or rotavirus infection showed that the virus-induced increase in JAK-STAT1/2 pathway activity could also be detected in this sample type. Influenza virus infection induced stronger immunity and this was associated with higher JAK-STAT1/2 pathway activity compared to rotavirus-infected patients (Figure 8).

**Figure 8:**
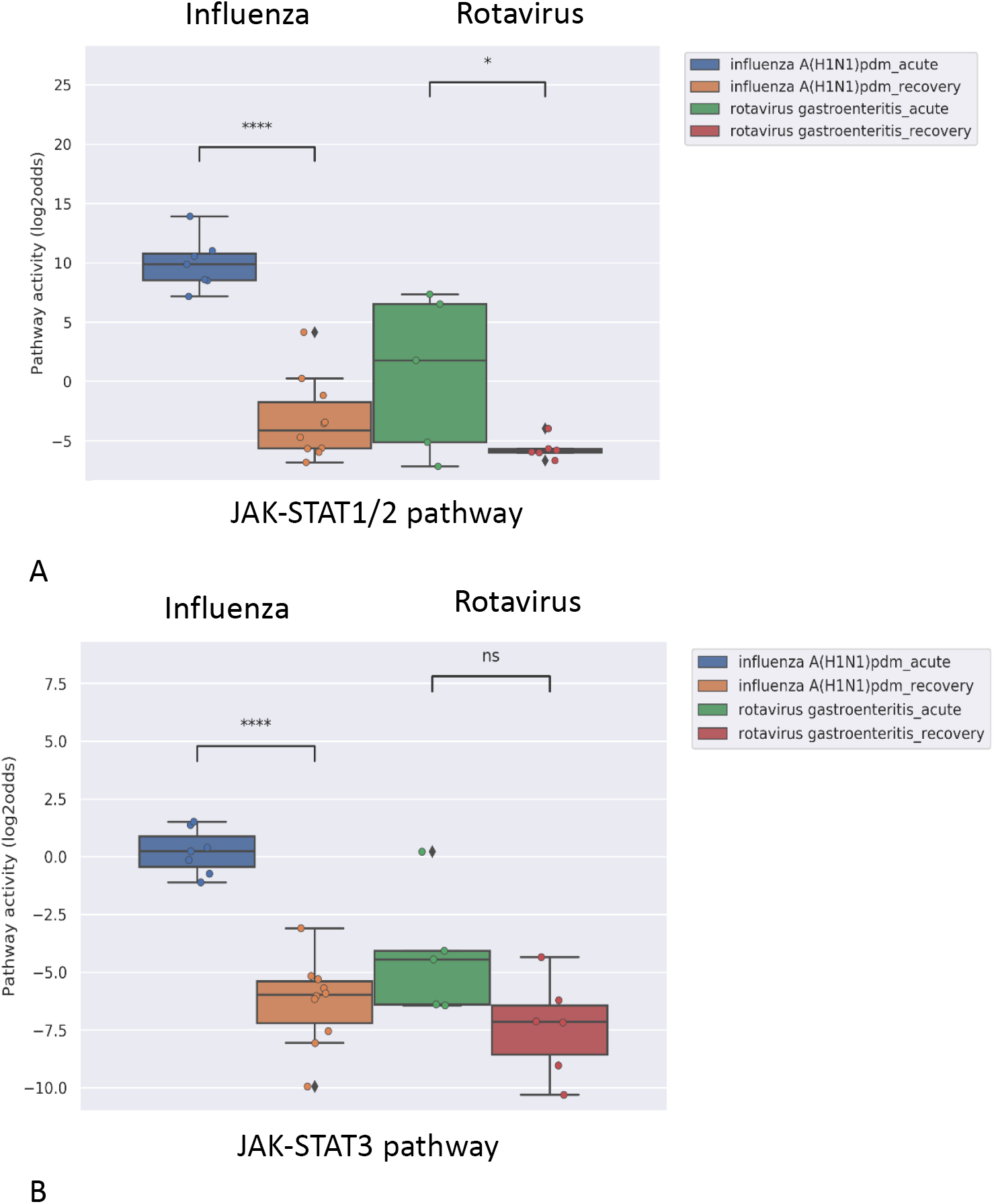
JAK-STAT1/2 (A) and JAK-STAT3 (B) pathway activity analysis of dataset GSE50628 (43). Influenza A(H1N1)pdm09 or rotavirus (gastroenteritis) infected patients. Whole blood samples were collected from patients on the day of admission in the acute phase (influenza, 1–3 days from disease onset; rotavirus, 2–4 days from disease onset and on the day of discharge) and recovery phase (Influenza, 4–9 days from disease onset; Rota, 7–11 days from disease onset), and were used for analysis. Pathway activity score on Y-axis in log2odds. Two sided t-test independent statistical tests were performed; p-values are indicated in the figures. P-values indicate: *p<0.05; ****p<0.0001.

#### Defining an upper threshold for JAK-STAT pathway activity in PBMC samples from healthy individuals

To enable clinical use of the JAK-STAT1/2 and JAK-STAT3 pathway tests to measure host immune response in virally infected patients for which no reference sample in healthy state is available, it would be of value to have a threshold value above which JAK-STAT pathway activity can be considered as increased. Two of the here analyzed datasets contained PBMC samples from healthy individuals (GSE34205 and GSE13486). Mean JAK-STAT1/2 pathway activity scores were comparable in the two independent datasets, consisting respectively of male (mean pathway activity score −5.84 log2odds, standard deviation SD 2.74) and female (mean −5.35 log2odds, SD 3.13) children and of adult (mean −6.50 log2odds, SD 2.56) individuals (Supplementary Table S3). Combining these groups, the mean JAK-STAT1/2 pathway activity score was −5.95 log2odds (SD 2.77) (Supplementary Table S3). Based on this, we propose to define an initial upper threshold for JAK-STAT1/2 pathway activity score in healthy individuals as the mean value + 2SD, resulting in a threshold of −0.41 log2odds for the pathway activity score, when measured on a PBMC sample. The upper threshold for JAK-STAT3 pathway activity score, calculated in the same way, was −4.42 log2odds, when measured in a PBMC sample.

## Discussion

We describe development of mRNA-based tests to quantify activity of the JAK-STAT1/2 and JAK-STAT3 signal transduction pathways in immune cells, and based on analysis of multiple clinical studies (influenza, RSV, yellow fever, dengue, rotavirus infections) provide evidence for value of the tests to quantify the host cellular immune response to a viral infection or vaccine.

### Value of measuring JAK-STAT pathway activity in viral infections

Measured JAK-STAT1/2 pathway activity scores in patient blood samples provided information on the adaptive immunity generated by viral infections, where a higher pathway activity was associated with development of a stronger adaptive immune response. This JAK-STAT pathway is known to perform key functions in generating an antiviral immune response (22),(44),(45). In most of the analyzed studies, PBMC samples had been collected that consist predominantly of lymphocytes and monocytes. Analysis of a Dengue infection and a Yellow Fever vaccination study, in which data were available from isolated T cells and monocytes, suggested that both lymphocytes and monocytes contributed to the increase in pathway activity seen in PBMCs - in line with the role of the JAK-STAT1/2 pathway in the immune response against viral infection. The increase in JAK-STAT1/2 pathway activity in PBMCs was only observed in viral infections, and not in bacterial infections. Interferons are well known to be specifically induced by a viral infection and to activate the JAK-STAT1/2 pathway to elicit an antiviral immune response (22),(44),(45). In line with this, the pathway test may have potential value in distinguishing between viral and bacterial infections.

Some viruses can induce a stronger and longer lasting adaptive immunity than others, and measured JAK-STAT1/2 pathway activity scores nicely reflected such differences. Influenza virus infection appeared to be associated with a higher JAK-STAT1/2 pathway activity in PBMCs than infection with RSV. Indeed, in general in the population the immune response generated by the RSV virus is lower and less persistent than immunity induced by the influenza virus (37). In Dengue infection, JAK-STAT1/2 pathway activity in PBMCs already started to decline within a few days after clinical presentation. This may have reflected the immunosuppressive effect of the Dengue virus, which has been reported to impair interferon-mediation induction of STAT1/2 activity (36),(46),(47). Finally, in the one study in which dendritic cells of patients with chronic persistent HBV infection were analyzed, JAK-STAT1/2 pathway activity was not increased at all. This was in line with described reduced functionality of DCs in chronic HBV infection, resulting in lack of adaptive immunity and chronic persistence of the HBV virus (38).

The immune response to infection with the same virus type is known to vary among patients (48). Overall we observed a large variation in JAK-STAT1/2 pathway activity between individual patients with a specific acute viral infection. While we cannot exclude that this may in part have been due to blood sampling at different time points during the disease, a more likely explanation is that it reflects differences in the individual patient capability to mount an adaptive immune response and generate immunity against the virus. Many factors influence this capability, like genetic variations, age, comorbidities like diabetes, and immune modifying drugs like corticosteroids (48),(49),(50).

Taken together, results suggest that measuring JAK-STAT1/2 pathway activity in PBMCs, or T cell subsets, and even in whole blood samples, during active viral infection may provide relevant information on the strength of the host immune response to induce adaptive immunity against the virus.

Changes in activity of the JAK-STAT3 pathway during viral infection frequently paralleled JAK-STAT1/2 pathway activity when measured on the same PBMC samples. However in contrast to the JAK-STAT1/2 pathway, JAK-STAT3 pathway activity increased more in monocytes than in T-cells. This suggests that activity of this signaling pathway is more associated with activity of the innate immune response, represented by the monocyte fraction in PBMC samples. Indeed, the JAK-STAT3 pathway is also involved in viral disease, and in addition to a role in generating a host immune response, its activity has been associated with severe inflammatory complications caused by a disbalance between adaptive and innate immune response, such as COVID-19 pneumonia (51),(52),(53),(54).

In two studies in which samples had been obtained from RSV and Dengue patients characterized by differences in disease severity, JAK-STAT3 pathway activity scores were higher in patients with a more severe disease course, although for Dengue only three such patient samples could be analyzed. Such high JAK-STAT3 pathway activity may have been induced by circulating cytokines such as IL6, IL10 or TNFα. These cytokines have been reported to be elevated in respiratory viral infections associated with severe infection of the lower airways and pneumonia, such as caused by RSV and more recently also SARS-CoV-2 (COVID-19), reflecting dominant activity of the innate immune response (55),(56),(57).

We believe that measuring JAK-STAT3 pathway activity in addition to JAK-STAT1/2 pathway activity may provide complementary information, that is, on the extent to which the innate immune response is activated, and potentially on the balance between adaptive and innate (inflammatory) response.

Absolute measured JAK-STAT pathway activity scores in healthy individuals were very comparable between different studies when obtained on the same type of blood sample. This allowed the calculation of an initial threshold value for PBMC samples above which a JAK-STAT1/2 or JAK-STAT3 pathway activity score may be considered as higher than normal, as found in healthy individuals. In view of the limited number of analyzed samples, these initial threshold levels need further confirmation. However, the high reproducibility between the different clinical studies suggests that defining a normal (“healthy”) range of JAK-STAT pathway activity in blood samples will be feasible. This will be of clinical relevance when assessing the immune response in virally infected patients, from whom usually no healthy reference blood sample will be available.

### Use of JAK-STAT pathway activity in vaccine development

Analysis of clinical vaccination studies provided support for the potential of using the JAK-STAT1/2 pathway activity score to predict vaccine immunogenicity and the induction of a cellular adaptive immune response. Yellow Fever (YF) vaccination induced a large increase in JAK-STAT1/2 pathway activity in PBMCs, both in T cells and monocytes, in all vaccinated individuals, within 3 days after vaccination. YF vaccine consists of an attenuated live virus, which infects dendritic cells, resulting in highly effective T cell activation, and excellent immunity (39). In contrast, after vaccination with a Trivalent Influenza Vaccine (TIV), no such increase in pathway activity was found, while during actual influenza virus infection JAK-STAT1/2 pathway activity did increase. This suggests that the vaccine had not induced adaptive cellular immunity, despite a satisfactory humoral antibody response in 22 out of 28 individuals (40). In contrast to the live virus YF vaccine, TIV contains inactivated virus. The inactivated virus vaccine may have lost the capability of the wild type virus to induce complete maturation and activation of dendritic cells, necessary to optimally activate T cells. Influenza vaccination not always induces satisfactory immunity and addition of an adjuvant compound to TIV was reported to increase immunogenicity (58),(59),(60). More recently, for this reason adjuvant-containing influenza vaccines have been introduced to increase influenza immunity in the aging population (61).

In addition to use for quantifying clinical effectiveness of a vaccine, the JAK-STAT1/2 pathway test may also be of value for *in vitro* vaccine development, as shown by our analysis of the study in which the immunogenicity of HIV had been investigated (62). HIV is not immunogenic by itself because the virus cannot infect DCs, however when infection was enabled *in vitro*, the DC activation process necessary to generate an immune response appeared to be fully functional (62). Only in the latter case, JAK-STAT1/2 pathway activity scores increased in DCs, together with increased activity of the NFκB pathway, together representing the main signaling pathways that need to be activated for effective DC antigen presentation (42).

The value of pathway activity measurement in immune cells is expected to be complementary to conventional IgG/IgM antibody testing (59). Because of the quantitative nature of the pathway tests, we expect that cellular effects of multiple vaccines can be rapidly quantitatively compared in vitro and a T cell immune response measured within 3-7 days after vaccination.

## Summary and conclusions

To the best of our knowledge, sofar it has not been possible to measure the cellular adaptive immune response in a blood sample from a patient. To further complete our signal transduction pathway testing platform, we have now developed tests to quantitatively measure functional activity of the JAK-STAT1/2 and JAK-STAT3 signal transduction pathways in different immune cell types, based on the principle of measuring and interpreting target gene mRNA levels of the signaling pathway transcription factor (6),(7),(8). We provide evidence that measurement of activity of the JAK-STAT1/2 signaling pathway in whole blood, PBMCs, or T-cell subsets, from patients with an infectious disease allows distinction between viral and bacterial infection and is informative on the strength of the adaptive cellular immune response against a viral infection. In addition, activity of the JAK-STAT3 pathway may be informative on the inflammatory component of the innate immune response and predictive of a more severe course of a viral infection, for example leading to a cytokine storm syndrome. Additional use of measuring JAK-STAT pathway activity in patient blood samples is expected to lie in prediction of response to specific immunomodulatory treatments, and monitoring effects of such treatments. Simultaneous measurement of activity of other signaling pathways, such as the NFκB, MAPK, PI3K, TGFβ and Notch pathways is possible on the same sample, and especially when performed on specific immune cell types, may allow more detailed information on the functionality of the immune response (63),(64),(65),(66) (unpublished results, manuscript in preparation). In addition signaling pathway analysis is expected to be useful to quantify immunogenicity and effectiveness of vaccines, especially with respect to the capacity to generate a T cell mediated adaptive immune response.

For clinical use in patients with a viral infection, it will be important to have the pathway activity test results in time to take clinical decisions. In the current study JAK-STAT pathway analysis was performed on Affymetrix expression microarray data. While of value for certain clinical studies, Affymetrix-based tests are not compatible with routine clinical use. For this reason, development of qPCR-based JAK-STAT pathway tests is in progress to reduce assay time (in principle within three hours) for routine clinical use, in a similar manner as described before for the ER signaling pathway test (11),(67).

## Relevance for the COVID-19 crisis

Rapid assessment of the cellular host immune response in a COVID-19 patient is needed to (a) improve prediction of prognosis and identification of patients at high risk for progression to COVID-19 pneumonia and ARDS, and prediction of clinical benefit of invasive ventilation at the ICU; (b) choose the optimal treatment, including immunomodulatory drugs; (c) assess effect of exploratory treatments on adaptive and innate (inflammatory) immune response; (d) develop better models to predict evolution of the pandemic; (e) accelerate vaccine development and clinical vaccine testing; (f) development of novel drugs to strengthen the adaptive immune response, while inhibiting the late inflammatory response associated with severe pneumonia and ARDS. We hope that the tests to quantitatively measure JAK-STAT1/2 and JAK-STAT3 signaling pathway activity in immune cells and blood samples can contribute to solutions for the COVID-19 crisis. Studies on measuring host immune response on blood samples from patients with COVID-19 infection are being initiated.

## Acknowledgement

We acknowledge Meng Dou and Henk van Ooijen for their valuable contributions to development of the JAK-STAT1/2 and JAK-STAT3 pathway models; and Diederick Keizer, Martijn Akse, and Sigi Neerken, for thoroughly reviewing the manuscript and providing valuable suggestions for improvement. We also wish to acknowledge all investigators who generated the used invaluable GEO datasets.

