## Supplementary information for "Quantitative measurement of activity of JAK-STAT signaling pathways in blood samples and immune cells to predict innate and adaptive cellular immune response to viral infection and accelerate vaccine development"

### Development of JAK/STAT1/2 and JAK-STAT3 pathway tests, Supplementary figures

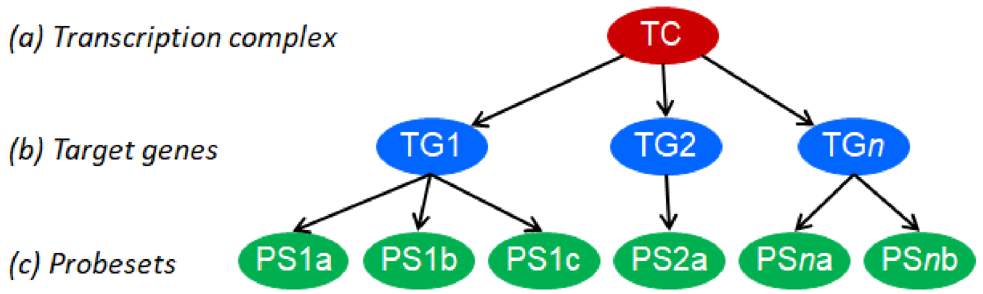


*Figure S1. Knowledge-based Bayesian computational pathway model. The Bayesian network structure used as a basis for our modeling approach shown as a simplified model of the transcriptional program of a cellular signal transduction pathway, consisting of three types of nodes: transcription factor, target gene, and microarray probe sets corresponding to target gene.*

*With permission (14).*

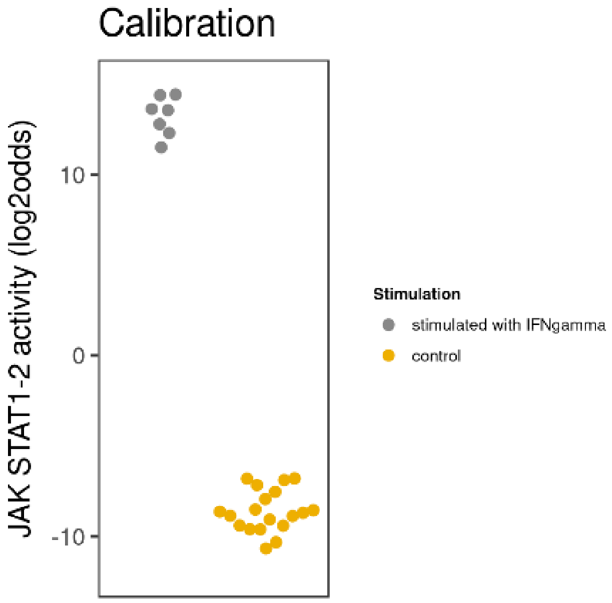

Calibration
JAK STAT1-2 activity (log2odds)
10
0
-10
Stimulation
stimulated with IFNgamma
control


**A**

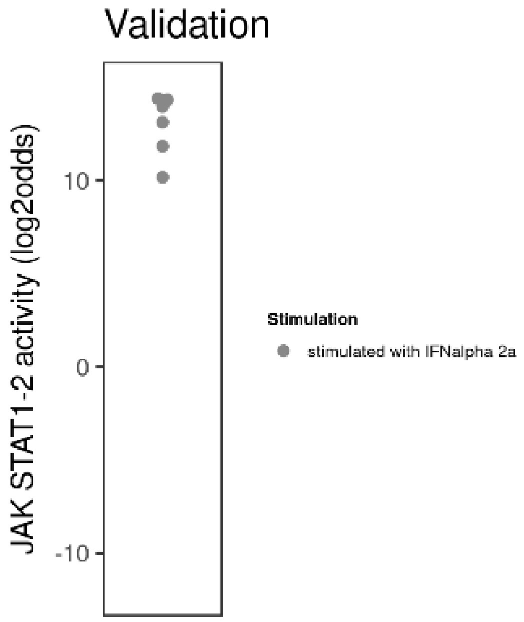

Validation
JAK STAT1-2 activity (log2odds)
10
0
-10
Stimulation
stimulated with IFNalpha 2a


**B**

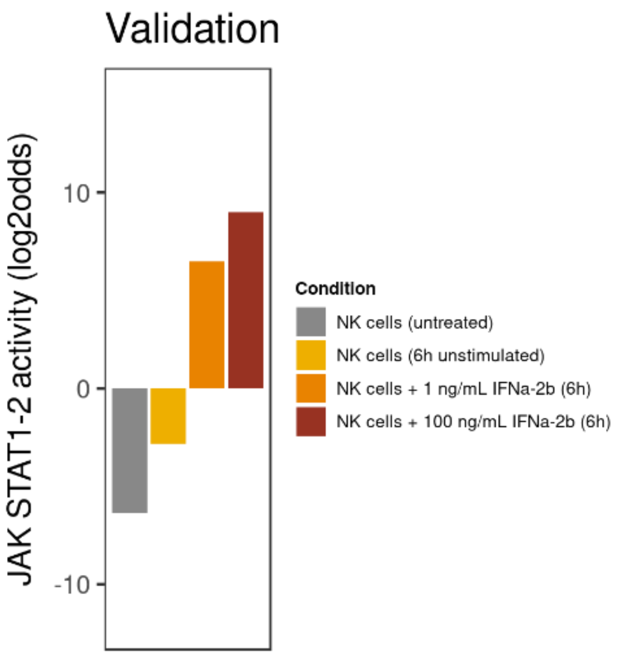
Validation
JAK STAT1-2 activity (log2odds)
10
0
-10
Condition
NK cells (untreated)
NK cells (6h unstimulated)
NK cells + 1 ng/mL IFNa-2b (6h)
NK cells + 100 ng/mL IFNa-2b (6h)

**C**

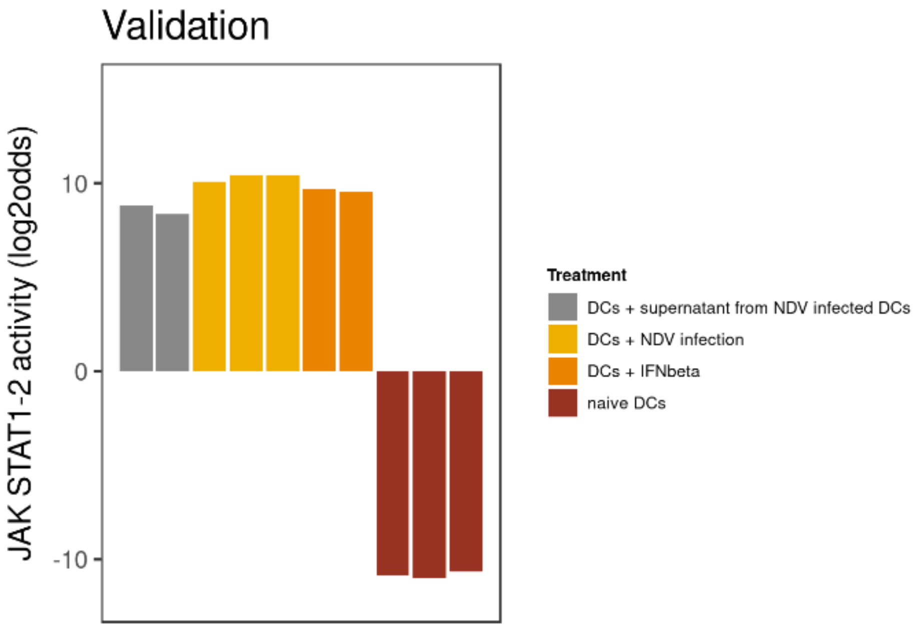
Validation
JAK STAT1-2 activity (log2odds)
10
0
-10
Treatment
DCs + supernatant from NDV infected DCs
DCs + NDV infection
DCs + IFNbeta
naive DCs

**D**

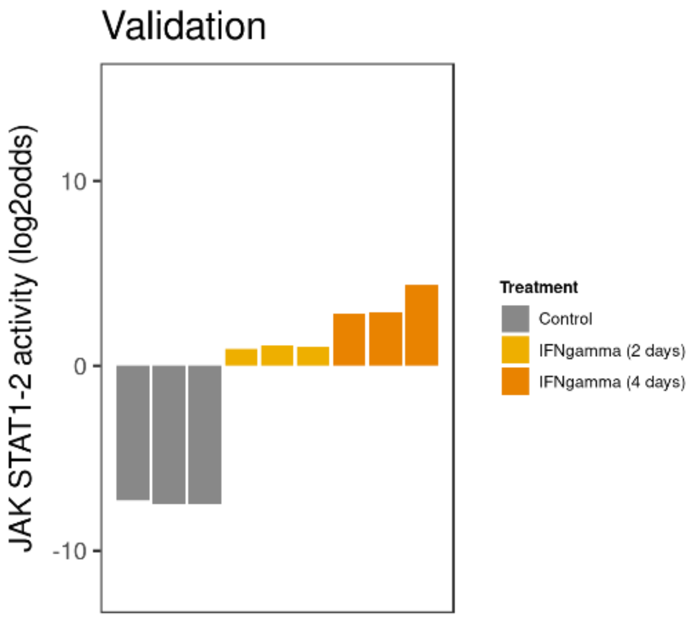
Validation
JAK STAT1-2 activity (log2odds)
10
0
-10
Treatment
Control
IFNgamma (2 days)
IFNgamma (4 days)

**E**

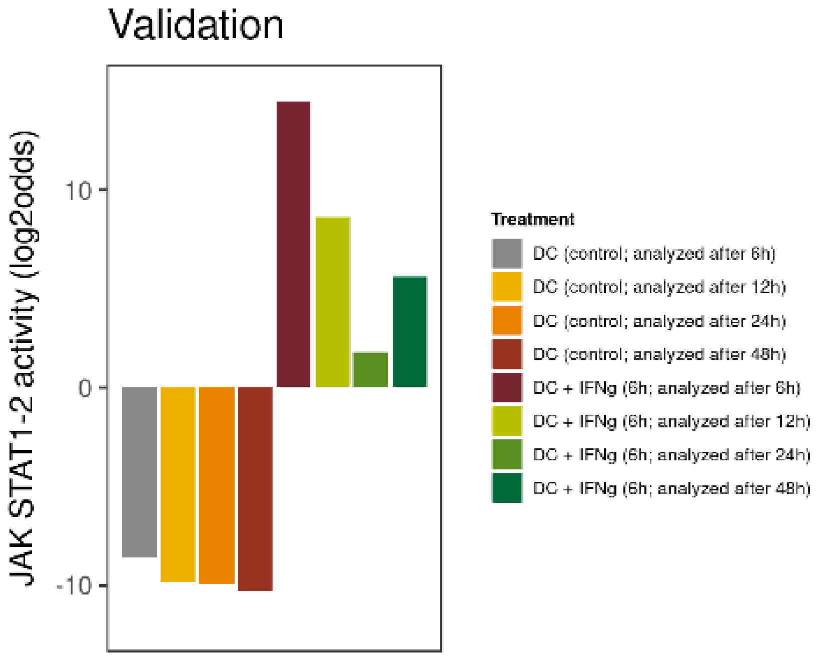
Validation
JAK STAT1-2 activity (log2odds)
10
0
-10
Treatment
DC (control; analyzed after 6h)
DC (control; analyzed after 12h)
DC (control; analyzed after 24h)
DC (control; analyzed after 48h)
DC + IFNg (6h; analyzed after 6h)
DC + IFNg (6h; analyzed after 12h)
DC + IFNg (6h; analyzed after 24h)
DC + IFNg (6h; analyzed after 48h)

**F**

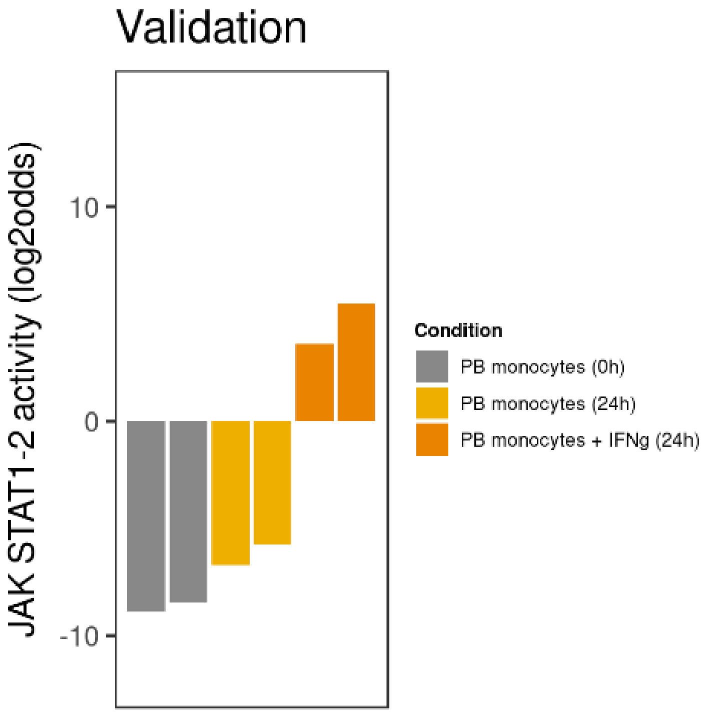


**G**

**Figure S2:** Calibration and biological validation of JAK-STAT 1/2 model.

**A:** GSE38351. Calibration of JAK-STAT1/2 pathway model with peripheral blood (PB) monocytes stimulated with 100 ng/mL interferon type II (IFNγ; n=7; active pathway) for 1.5 hours and unstimulated n=19; inactive pathway).

**B-D**. Validation of JAK-STAT1/2 pathway model for IFN type I induced pathway activity.

**B**. GSE38351. Peripheral blood monocytes stimulated for 1.5 hrs with IFNα2a (n=7).

**C.** GSE15743. Peripheral natural killer (NK) cells, untreated, cultured for 6 hours without stimulation, treated for 6 hours with 1 ng/mL recombinant IFNα-2b (low interferon α) and treated with 100 ng/mL recombinant IFNα-2b (high interferon α).

**D**. GSE52081. Dendritic cells, exposed to supernatant of Newcastle Disease Virus (NDV)-infected cells, infected with NDV, treated with IFNβ or kept as naïve dendritic cells.

**E-G:** Validation of the JAK-STAT1/2 pathway model for IFN type II-induced pathway activity

**E:** GSE58096.THP1 human monocyte-like cells were untreated or treated with IFNγ for 2 or 4 days.

**F:** GSE11327. Immature monocyte-derived dendritic cells were untreated or matured with 30 ng/mL lipopolysaccharide (LPS) and 1000 U/mL IFNγ for 6 hours, and analyzed at 6, 12, 24 and 48 hours. Each grey (treated) bar matches with the respective coloured bar (untreated).

**G:** GSE11864. Peripheral blood monocytes (from 2 donors) were directly analyzed or cultured in the absence or presence of 100 U/mL IFNγ for 24 hours.

*The pathway activity score is presented on a log2 odds scale. Two-sided Wilcoxon signed-rank statistical tests were performed, p-values are indicated in de figures. In case fewer than 4 samples needed presentation, bar plots are used instead of dot blots.*

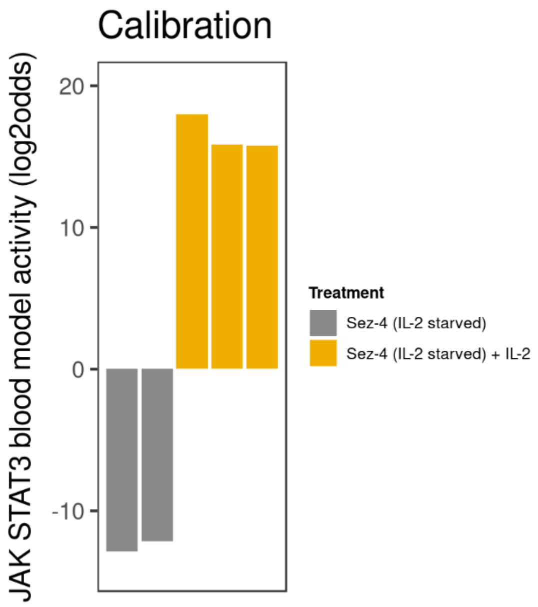


**A**

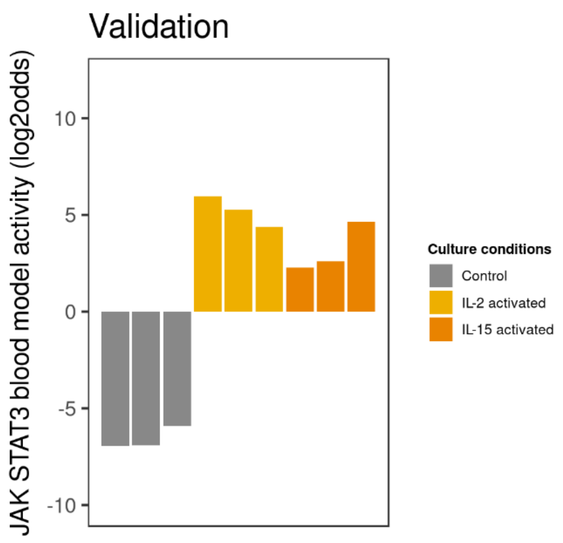
Validation
JAK STAT3 blood model activity (log2odds)
10
5
0
-5
-10
Culture conditions
Control
IL-2 activated
IL-15 activated

**B**

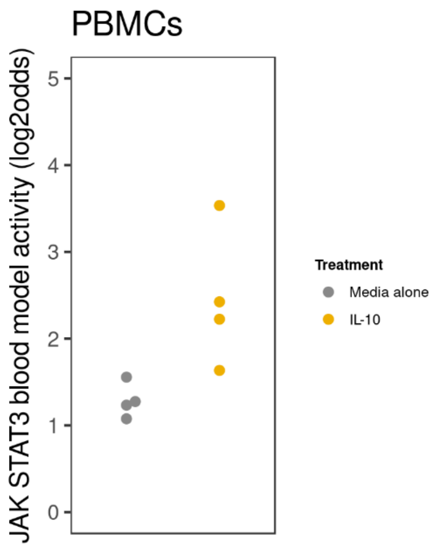
PBMCs
JAK STAT3 blood model activity (log2odds)
5
4
3
2
1
0
Treatment
Media alone
IL-10

**C**

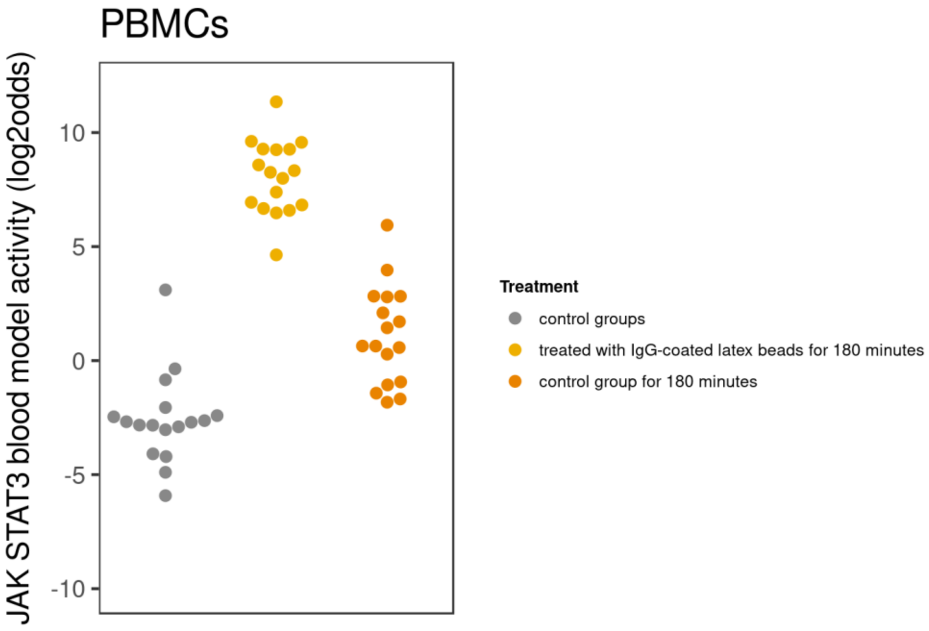
PBMCs
JAK STAT3 blood model activity (log2odds)
10
5
0
-5
-10
Treatment
control groups
treated with IgG-coated latex beads for 180 minutes
control group for 180 minutes

**D**

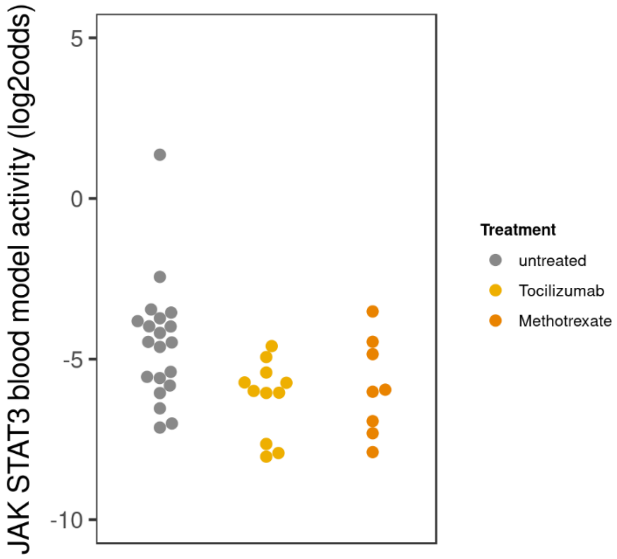
JAK STAT3 blood model activity (log2odds)
5
0
-5
-10
Treatment
untreated
Tocilizumab
Methotrexate

**E**

Figure S3. Calibration and biological validation of JAK-STAT3 pathway model for use on blood cells (JAK-STAT3-B model).

**A:** GSE8687. Calibration of the JAK-STAT3-B pathway model with CD4+ T cells derived from leukemic (Sezary) cells of a patient with a cutaneous T-cell lymphoma. Cells were starved of Interleukin-2 (IL-2; n=2; inactive pathway) or cultured with IL-2 (n=3; active pathway).
**B:** GSE8685. Validation of the JAK-STAT3-B model on independent samples from Sez-4 T cell lymphoma cells that were cultured 16 hours in absence of IL-2 (IL-2 starved) and subsequently treated with 200 U IL-2, 20 ng/mL IL-15, 100 ng/mL IL-21 or vehicle for 4 hours.
**C**. GSE43700. PBMCs from blood of healthy human donors (n=4), stimulated by IL-10 (R&D Systems) 10ng/ml or vehicle for 24 hours.
**D**. GSE8507. PBMCSs were measured directly, or stimulated with IgG-coated latex beads or control for 180 minutes.
**E**. GSE45867. Knee synovial biopsies were taken from patients with RA, before (untreated) and after 12 weeks of treatment with Tocilizumab (IL6 inhibitor) or methotrexate (treatment naive pts). Tocilizumab was clinically more effective. Paired patient samples, analyzed with paired T-test: comparison between untreated and Tocilizumab (n=12), p=0.005; between untreated and methotrexate (n=8): p=0.131 (ns).
*The pathway activity score is presented on a log2 odds scale. Two-sided Wilcoxon signed-rank statistical tests were performed unless otherwise indicated, p-values are indicated in de figures. In case fewer than 4 samples needed presentation, bar plots are used instead of dot blots.*

### Supplementary Methods

**Selection of target genes for Bayesian models for JAK-STAT1/2 and JAK-STAT3 pathway models**
For each putative target gene of a pathway-associated transcription factor, evidence was assessed for the presence of a binding element in the gene promoter region, functionality of the binding element (e.g., in promoter-luciferase experiments), binding of the transcription factor to the respective response/enhancer element *in vivo* (e.g., using ChIPseq) and/or *in vitro* (using Electrophoretic Mobility Shift Assay, EMSA), and differential expression with pathway activation. Gene selection was also based on consistency of evidence as reported by multiple research groups for multiple cell/tissue types. Around 20 genes per pathway were selected, which is high enough to enable robustness and sensitivity of the pathway assay, while allowing for maximal

specificity. Because the target genes only function to read out transcription factor activity, they were selected based on evidence for reproducible and specific transcription factor-induced transactivation across various cell types, and not based on function of encoded proteins. Probesets on the Affymetrix HG-U133Plus2.0 microarray associated with the target genes were selected based on the Bioconductor package available in R and manual curation using the latest information available on the UCSC Genome Browser (www.genome.ucsc.edu) (1).

### Supplementary Tables

**Table S1. List with target genes used in the Bayesian models to develop JAK-STAT1/2 and JAK-STAT3 signal transduction pathway tests**

JAK-STAT1/2 signaling pathway: BID, GNAZ, IRF1, IRF7, IRF8, IRF9, LGALS1, NCF4, NFAM1, OAS1, PDCD1, RAB36, RBX1, RFPL3, SAMM50, SMARCB1, SSTR3, ST13, STAT1, TRMT1, UFD1L, USP18, ZNRF3.

JAK-STAT3 signaling pathway: AKT1, BCL2, BCL2L1, BIRC5, CCND1, CD274, CDKN1A, CRP, FGF2, FOS, FSCN1, FSCN2, FSCN3, HIF1A, HSP90AA1, HSP90AB1, HSP90B1, HSPA1A, HSPA1B, ICAM1, IFNG, IL10, JUNB, MCL1, MMP1, MMP3, MMP9, MUC1, MYC, NOS2, POU2F1, PTGS2, SAA1, STAT1, TIMP1, TNFRSF1B, TWIST1, VIM, ZEB1.

**Table S2. References for target gene selection**

JAK-STAT1/2 pathway
(2),(3),(4), (5),(6),(7),(8),(9),(10),(11),(12)
JAK-STAT3 pathway
(13),(14),(15),(14),(16)

**Table S3 JAK-STAT pathway activity scores in PBMC samples of health individuals**

S3A. Healthy individual samples from datasets GSE34205 and GSE13486, separate groups.

| GSE dataset | Female (F) /make (M) | JAK-STAT1/2 pathway activity score *(log2odds)* Mean | JAK-STAT1/2 pathway activity score *(log2odds)* *Standard deviation* | JAK-STAT3 pathway activity score *(log2odds)* *Mean* | JAK-STAT3 pathway activity score *(log2odds)* *Standard deviation* |
|---|---|---|---|---|---|
| GSE34205 | F | -5.35 | 3.13 | -7.49 | 1.15 |
| GSE34205 | M | -5.84 | 2.74 | -8.18 | 1.91 |
| GSE13486 | | -6.50 | 2.56 | -7.73 | 1.79 |

S3B. Healthy individual samples from datasets GSE34205 and GSE13486 were combined.

| GSE dataset | Female(F) /male (M) | JAK-STAT1/2 pathway activity score *(log2odds)* *Mean* | JAK-STAT1/2 pathway activity score *(log2odds)* *Standard deviation* | JAK-STAT3 pathway activity score *(log2odds)* *Mean* | JAK-STAT13 pathway activity score *(log2odds)* *Standard deviation* |
|---|---|---|---|---|---|
| GSE34205 GSE13486 | F/M | -5.95 | 2.77 | -7.86 | 1.72 |
